# *Mycobacterium tuberculosis* Rv3194c efficiently facilitates mycobacterial-lung epithelial interaction through its HA-binding site

**DOI:** 10.1101/2020.02.24.963918

**Authors:** Dongyue Zhao, Chen Xu, Danfeng Lin

## Abstract

*Mycobacterium tuberculosis* adhesins are surface-exposed molecules that mediate pathogen-host interaction, a fundamental step towards host infection. Here we show that serine protease (Rv3194c) promotes mycobacterial infection to lung epithelial through its hyaluronic acid (HA)-binding site. Both enzyme-linked immunosorbent assay and surface plasmon resonance analysis revealed that Rv3194c bound to HA. Utilizing synthetic peptides, we next defined HA-binding site of 20 amino acids from 91 to 110 of Rv3194c (P91-110). Immunofluorescence assay and an FACScan showed that Rv3194c was interacted with A549 cells (human lung epithelial cells), and its interaction was abolished by the addition of hyaluronidase or P91-110. Experimental infection *in Vitro* revealed that Rv3194c participates in attachment of recombinant *Mycobacterium smegmatis* (Rv3194c/MS) to A549 cells, and P91-110 treatment of A549 cells almost inhibited Rv3194c/MS-A549 cells interaction. To provide *in vivo* evidence, we constructed a reporter strain of *M. smegmatis* expressed a derivative of the firefly luciferase that is shifted to red (FFlucRT) in combination with Rv3194c (Rv3194c+FFlucRT/MS) to infect the rodents and monitor the progression of the disease. Using bioluminescence imaging and bacterial counts in lung tissue confirmed that Rv3194c dramatically enhanced the persistence of *M. smegmatis*. In addition, treatment of intratracheal Rv3194c+FFlucRT/MS-infected mice with P91-110 significantly suppressed the growth of Rv3194c+FFlucRT/MS *in vivo*. Taken together, these results demonstrate that Rv3194c was identified as a HA-binding adhesin, and P91-110 as anti-adhesion agents has potential for therapeutic and prophylactic interventions in mycobacterial infection.

---

*Mycobacterium tuberculosis* to stick to the host cell is the most important step for the pathogen to start the infection (1). During the infection process, it has been clearly shown that *M. tuberculosis* interacts with host cells and enters them (2). Such cells include epithelial cell, macrophage and dendritic cell (2). This process of attachment and subsequent entry into host cells is critical to facilitate delivery of toxin and virulence factors to the host cell (3), helps the *M. tuberculosis* to maintain their position and resist the host immunity (4). Mycobacterial adhesins, located on the bacterial surface or on pili, specifically interact with extracellular matrix (ECM) components and mediate *M. tuberculosis-*host cells interaction (5). The ECM components include collagen, bone sialoprotein, fibronectin, elastin, laminin, glycosaminoglycans (GAGs) and vitronectin (5). Because attachment of *M. tuberculosis* to host cell confers tissue tropism (6), they may carry adhesins for more than one target surface and may employ more than one adhesin for binding to a substrate. In analyzing the process of attachment, it was found that a variety of adhesins were necessary for the infection of *M. tuberculosis* with the lung. Pili are a surface structure similar to hair, and their distal tip contains an adhesin, Curli Pili encoded by *Rv3312A*, which binds to laminin on macrophages and epithelial cells surface (7, 8). However, *M. tuberculosis* has also been found to contain a large of adhesins that do not involve pili. The Pro-Glu (PE) polymorphic GC-rich repetitive sequence (PGRS) protein encoded by *Rv1818c* participates in attachment of *M. tuberculosis* to epithelial cells or macrophages through binding to fibronectin (9). Early secreted antigen ESAT-6 encoded by *Rv3875* play a role in dissemination of *M. tuberculosis* via binding to laminin on human lung epithelial cells (10). *M. tuberculosis* Cpn60.2 encoded by *Rv0440* binds to CD43 and effectively facilitates attachment of *M. tuberculosis* to macrophages (11). *Mycobacterium* cell entry-1 protein encoded by *Rv0169* participates in attachment and subsequent internalization of *M. tuberculosis* to epithelial cells (12). The membrane protein encoded by *Rv2599* binds to collagen, fibronectin and laminin; N-acetylmuramoyl-L-alanine amidase encoded by *Rv3717* binds to fibronectin and laminin; L, D-transpeptidase encoded by *Rv0309* binds to fibronectin and laminin (13). The malate synthase encoded by *Rv1837c* has adapted to function as an ashesin which binds to laminin and fibronectin on lung epithelial cells surface (14). PE-PGRS family protein Wag22 encoded by *Rv1759c* (15), antigen 85 complex encoded by *Rv0129c and Rv1886c* (16) and glutamine synthetase A1 encoded by *Rv2220* (17) respectively bind to fibronectin. PstS-1 (the *M. tuberculosis* 38 kDa glycoprotein) encoded by *Rv0934* (18) and LpqH glycolipoprotein (the 19 kDa antigen) encoded by *Rv3763* (19) act as an adhesin that can bind the mannose receptor and promote the phagocytosis of the mycobacteria. Extracellular Mycobacterial DNA binding protein 1 (MDP-1) promotes mycobacterial-lung epithelial interaction through binding HA (20). The heparin-binding hemagglutinin (HBHA) encoded by *Rv0475* binds to heparin and involves in the adherence to lung epithelial cells (21). With the emergence of multi-resistant mycobacteria and the mixed infection with the virus, the treatment of Tuberculosis (TB) is becoming more and more difficult (22). Currently, the anti-adhesion therapy that interferes with the ability of the *M. tuberculosis* to adhere to tissues of the host at the early stages of infection is good therapeutic strategies (23). Adhesins of *M. tuberculosis* initiate the process of adhesion between host and bacteria. The adhesion process is not determined by a single or a few adhesins. Therefore, to explore new *M. tuberculosis* adhesins in the research process is essential for understanding of the pathogenesis of *M. tuberculosis* and discovery new candidate drug against TB.

The interaction with the epithelia is considered to be critical to enable the bacteria to disseminate and dominate the outcome of the disease (24). In addition to the adhesins described above, *M. tuberculosis* may have the other adhesins. Analysis of Rv3194c using the SMART tool (http://smart.embl-heidelberg.de/) has shown that 123-192 amino acids are a PDZ domain, 131-194 are an SCOP (Structure Classification of Proteins, http://scop.mrc-lmb.cam.ac.uk/scop/data/scop.b.html) domain, d1g9oa_, of which the members might possess the capacity of binding DNA, ATP or proteins associated with apoptosis and 232-330 are a Lon C domain. In addition, the minimal HA-binding requirement (_**94**_**R**DLVYPPG**K**_**102**_), the B(X_7_)B motif (25), was found in the sequence of Rv3194c, which might possess adhesion characteristics. The recent study (26) demonstrated that Lon C domain in Rv3194c hydrolyzes BSA, casein, gelatin, and milk, and degraded the complement components, C3b and C5a. However, the substrates recognized by PDZ domain have been unidentified, and whether its B(X_7_)B (25) motif possesses adhesion characteristics has not been known. Therefore, our works focus on the adhesion role of the extracellular Rv3194c. The results indicate that Rv3194c mediates the mycobacteria-lung epithelial interactions through binding of peptide corresponding to an amino acid sequence of Rv3194c at the 91-110 position (P91-110), HA-binding site, to HA in cell surface. In addition, the growth of recombinant *M. smegmatis* (rMs) *in vivo* was notably reduced when the mice was treated with P91-110. In summary, the Rv3194c acts as a novel adhesin, and its peptide, P91-110, may be a candidate drug against TB.

## Results

### Expression and purification of the Rv3194c protein

To obtain the potential structural information of the Rv3194c protein, we first examined its sequence with several types of structure prediction software. TMHMM server (http://www.cbs.dtu.dk/services/TMHMM-2.0/) predicted a transmembrane helix from 1 to 27. Because the hydrophobic sequence from the trans-membrane domain might make the protein more difficult to express, *Rv3194c* without the trans-membrane helix sequence was cloned into a His-tag vector to produce a His-tagged fusion protein. To facilitate the activity of the target protein and to increase the amount of the soluble fraction, the expression condition with 0.5 mM IPTG at 23°C for 12 h was optimized. After ultra-sonication, the soluble protein in the supernatant was purified by affinity chromatography with Ni^2+^. After purification, the recombinant Rv3194c showed the expected molecular size (35 kDa) when analyzed using SDS-PAGE (**Fig. 1A**). A western blotting further revealed that an approximately 35 kDa protein band was the Rv3194c protein after it was probed with the anti-Rv3194c antisera (**Fig. 1B**). The 35 kDa protein from cell lysate of BCG, as the positive control, was recognized by anti-Rv3194c antisera (**Fig. 1B**). Followed by using the Superdex 75 gel chromatography methods for the protein’s separation and purification, we have got the more than 99% protein of Rv3194c (**Fig. 1C**). Finally, mass spectrometry and peptide mass fingerprinting results confirmed that the expected protein, Rv3194c (Accession No. NP_217710.1), was obtained from H37Rv (**Fig. 1D**).

**FIG 1.**
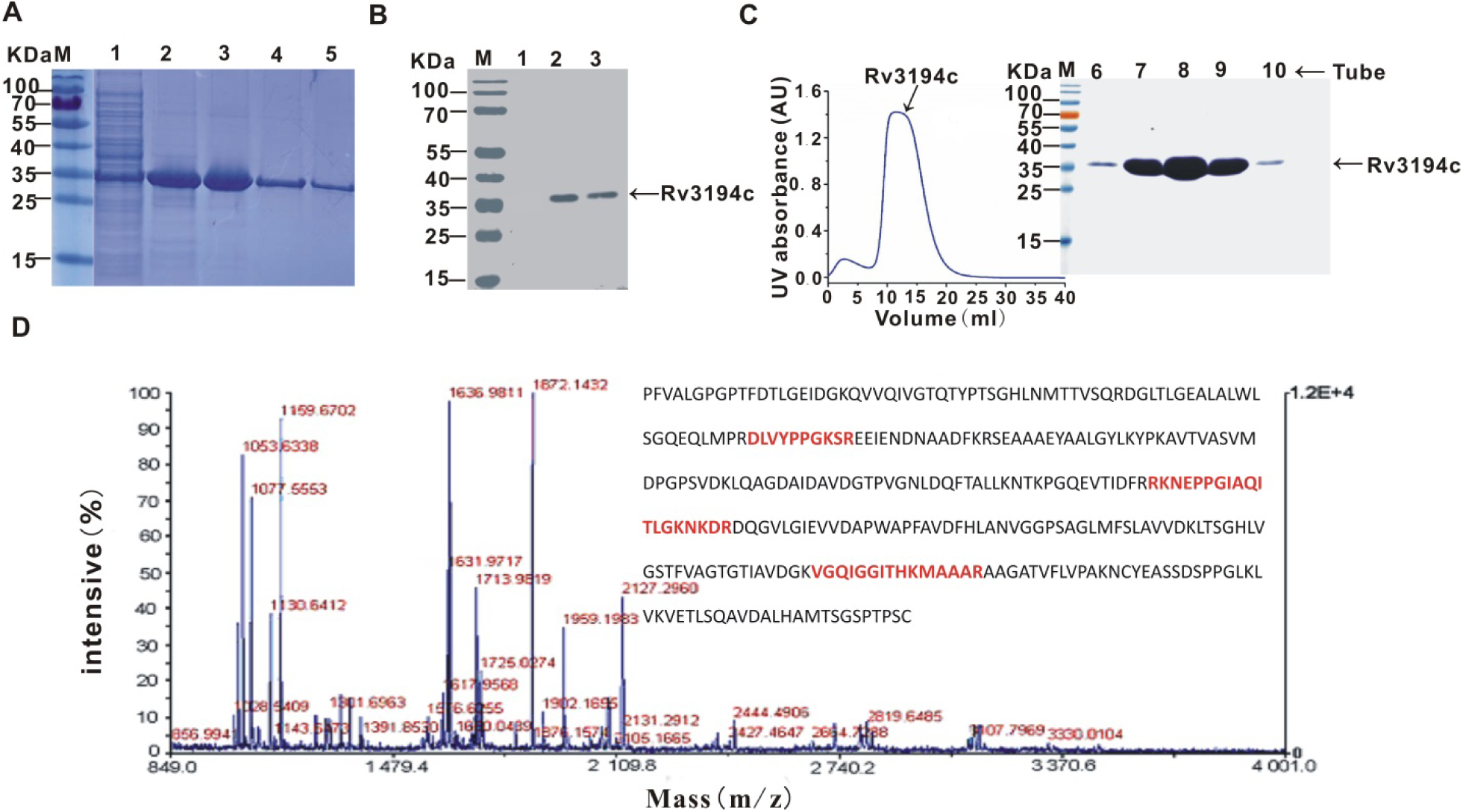
Expression, purification and validation of the Rv3194c protein. **A**, Analysis of Rv3194c protein by SDS-PAGE. Lane 1, the supernatant of induced cells after disruption; Lane 2, agarose resin after supernatant to pass through column; Lane 3, agarose resin after column washed by wash buffer; Lane 4, purified Rv3194c protein was eluted by 20 mM Tris, 200 mM imidazole and 150 mM NaCl, (pH 7.5); Lane 5, purified Rv3194c protein was eluted using 20 mM Tris, 500 mM imidazole and 150 mM NaCl, (pH 7.5). **B**, Analysis of Rv3194c protein by western blotting. Lane 1, pET28a-Rv3194c without induction served as a negative control; Lane 2, pET28a-Rv3194c with induction; Lane 3, the cell lysate of BCG served as a positive control. **C**, Gel filtration chromatography of the Rv3194c protein. Proteins collected in different tube from 6 to10 were respectively detected by SDS-PAGE. D MALDI-TOF peptide mass fingerprint spectrometry for the recombinant Rv3194c. The sequence of Rv3194c covering these fragments is displayed as bolded in red.

### Determination of the HA-binding Site of Rv3194c

Analysis of Rv3194c revealed that the minimal HA-binding requirement (_**94**_**R**DLVYPPG**K**_**102**_), the B(X_7_)B motif (25), was found in the sequence of Rv3194c, which might possess adhesion characteristics. To elucidate this hypothesis, we examined whether Rv3194c binds to HA. The ELISA showed that Rv3194c bound to HA, but not mannose, dextran and galactose (**Fig. 2A**); the Rv3295, as a negative control protein, does not bind to HA, dextran, mannose and galactose (**Fig. 2A**). To explore the role of extracellularly occurring Rv3194c, HA-Sepharose Chromatography was used to examine the binding capacity of Rv3194c to HA. HA-binding protein, Rv3194c protein on western blotting analysis, was eluted by PBS containing 1.5 M NaCl (**Fig. 2B**). To determine its binding site, we examined the HA-binding activity of 20-mer of synthetic peptides covering the entire sequence of Rv3194c. ELISA showed that the HA bound to the peptide corresponding to an amino acid sequence of Rv3194c at the 91-110 position (P91-110) (**Fig. 2C**). The results indicated that Rv3194c bound to HA through its P91-110.

**FIG 2.**
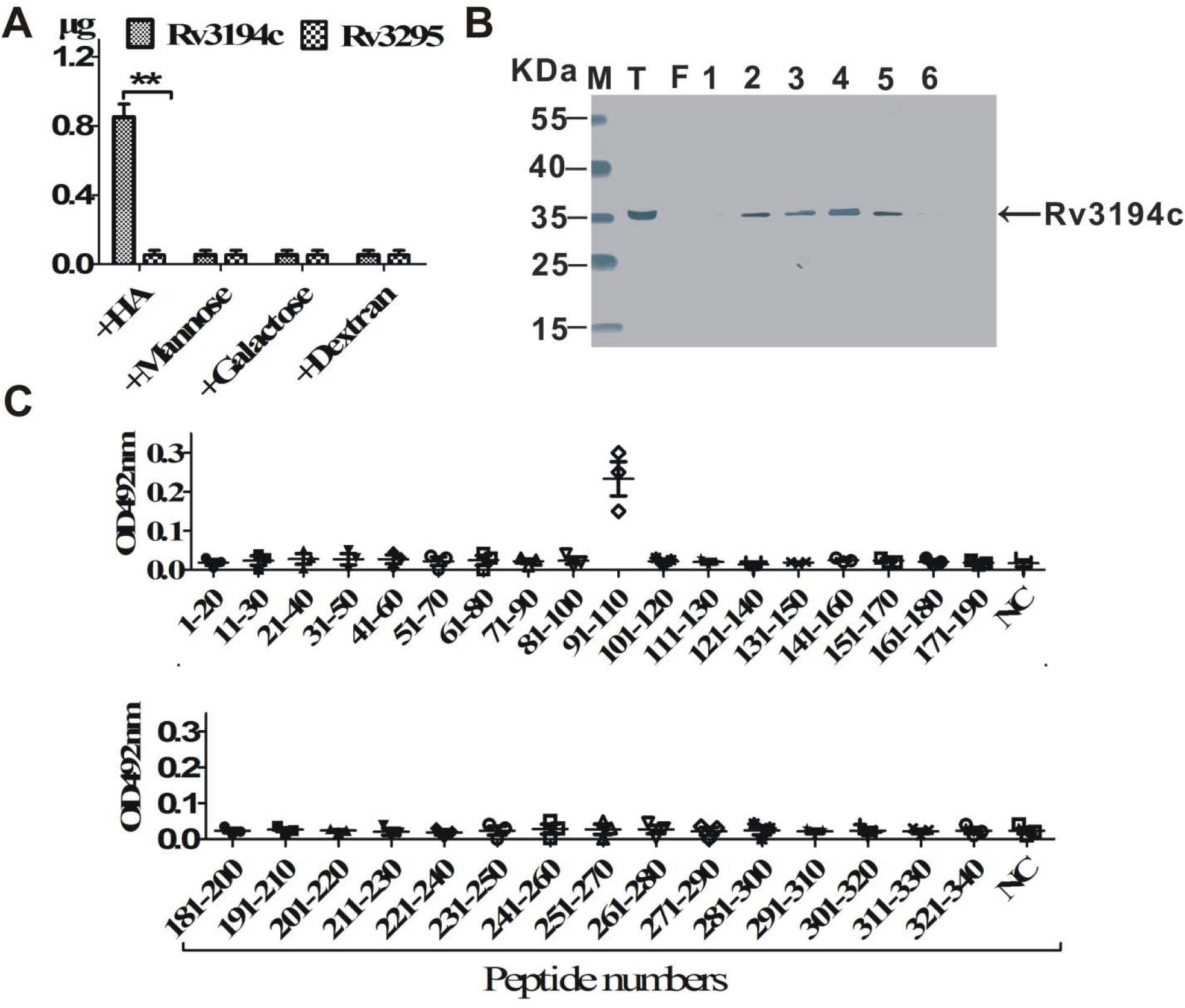
Rv3194c binds to HA. **A**, The content of Rv3194c in the immobilized wells was tested by ELISA. Rv3295 instead of Rv3194c served as a negative control. **B**, Analysis of HA-binding proteins using HA-Sepharose chromatography. T, total Rv3194c protein applied to chromatography; F, flow-through; Lane1-6, Rv3194c eluted by 1.5M NaCl. **C**, The Rv3194c-specific HA binding region was determined by ELISA. 20-mer of sequential peptides corresponding to the amino acid sequence of Rv3194c was synthesized. *OD*, optical density; *NC*, negative control without peptide.

### SPR analysis of the interaction between Rv3194c and the HA

Based on results as described above, Rv3194c and P91-110 were respectively treated as an immobilized ligand in the following studies. To elucidate the molecular interaction, an SPR analysis was performed. Rv3194c, P91-110 and Rv3295 were immobilized on the CM5 sensor chip until they had gained 3253, 3212 and 3034 resonance units (RUs). HA (10 μg/mL) was injected into the immobilized cells with either Rv3194c or P91-110 for 30 s. As predicted from the analysis by HA-Sepharose Chromatography, the interaction of Rv3194c or P91-110 with HA was observed until they had gained 85 RU or 110 RU, respectively (**Fig. 3A**). HA had no interactions with Rv3295 or the empty sensor (data not shown). The affinity (*K*_*D*_) of HA bound to Rv3194c or P91-110 was 2×10^−8^ mol/L or 1.6×10^−8^ mol/L, respectively (**Fig. 3B**). The reason for the difference of *K*_*D*_ value, a part of the binding site of synthetic peptide may be masked during immobilization procedure. In contrast, different types of carbohydrates including mannose, dextrose and galactose, lacked the ability to bind to Rv3194c or Rv3295 (data not shown). The results of the SPR analysis indicate that Rv3194c bound to HA with highest affinity, and the P91-110, HA-binding Site, was also further confirmed (**Fig. 3**).

**FIG 3.**
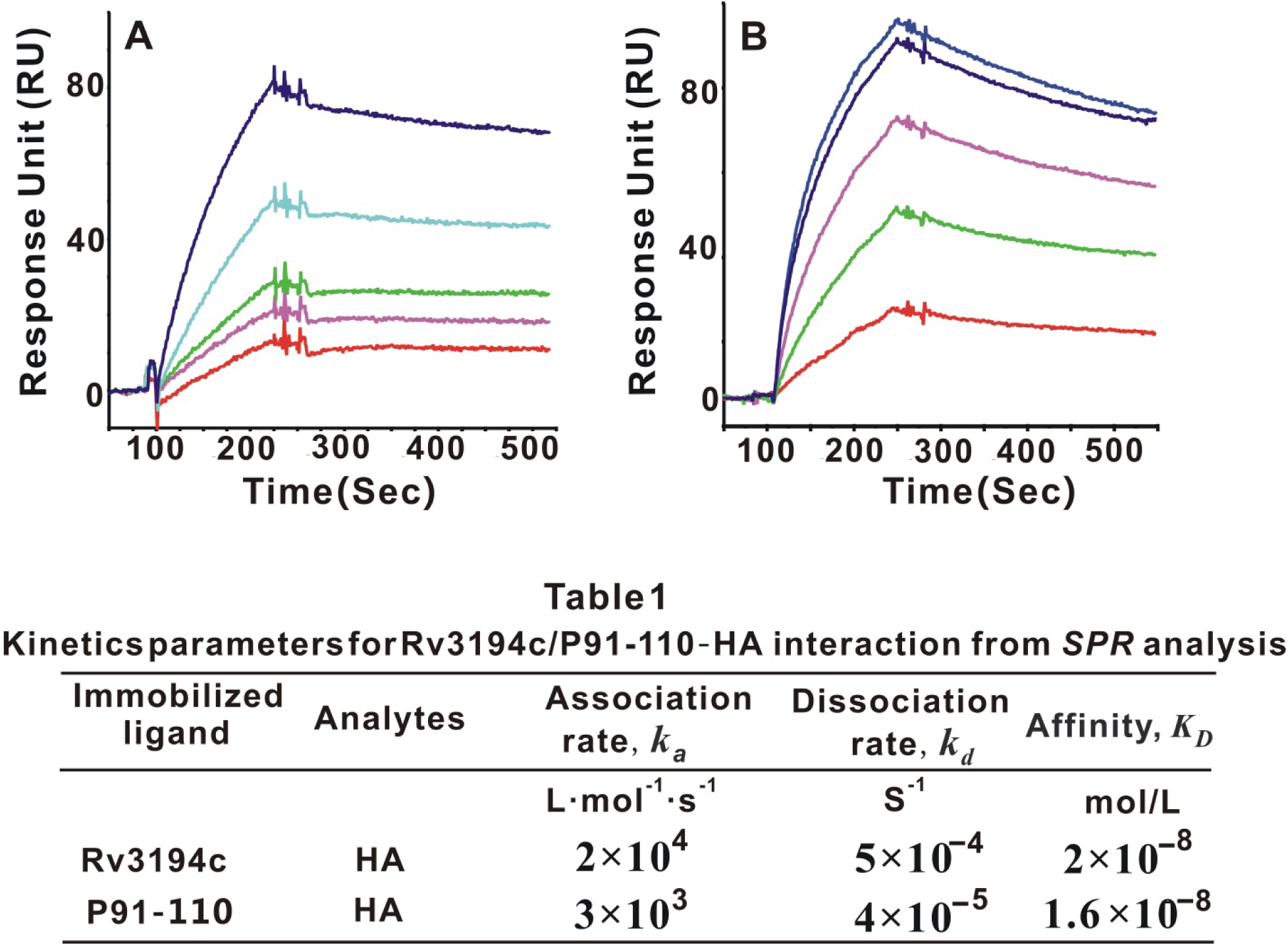
The interaction of Rv3194c/P91-110 with HA analyzed by surface plasmon resonance. Results for Rv3194c (**A**) and P91-110 (**B**) are shown. Injections of different concentrations of HA flowed over a surface chip at a flow rate of 10 μL/min. Concentration of the analyte: 15, 30, 60, 120 and 240 nM.

### Rv3194c Binds to the A549 Human Lung Epithelial Cells

HA, a glycosaminoglycans (GAGs), located in the surface of Lung Epithelial Cells, possessing diverse biological functions (27). The results (**Figs. 2 and 3**) prompted us to examine the possibility that the Rv3194c may participate in the binding to A549 epithelial cells. Rv3194c and P91-110 conjugated with red fluorescent group known as Cy3 at its N-terminal (Cy3-P91-110) were respectively incubated with A549 cells to determine if they can bind to the epithelial cells. Confocal laser microscopy showed that Rv3194c and Cy3-P91-110 attached to the cells, but not that of Rv3295 (**Fig. 4A**). The binding of Cy3-P91-110 to A549 cells was obvious by 10 min, and by 1 hour after the addition, the binding had reached a plateau; More than 98% of the A549 cells were Cy3-P91-110-positive 1 h later (**Fig. 4B**). In contrast, binding of Rv3194c was delayed, and 20% of cells were Rv3194c negative even at the plateau phase, suggesting that the binding of P91-110 to cells is more efficient than that of Rv3194c (**Fig. 4B**). To determine whether the binding of Rv3194c to A549 cell relies on HA, the subsequent experiments were conducted. We studied the ability of the exogenous hyaluronidase, HA or P91-110 to inhibit the reaction. FACScan showed that the percentage of binding of Rv3194c to untreated A549 cells was more than 98% (**Fig. 4C**); percentage of binding of Rv3194c to A549 cells pretreated with hyaluronidase, HA or P91-110 was respectively less than 5.2%, 51% or 3.2% (**Fig. 4C**), indicating that addition of exogenous hyaluronidase or P91-110 abolished Rv3194c-A549 cells interaction compared to that of added HA. In total, these findings indicated that Rv3194c bound to epithelial cells through its P91-110 interaction with HA in cells.

**FIG 4.**
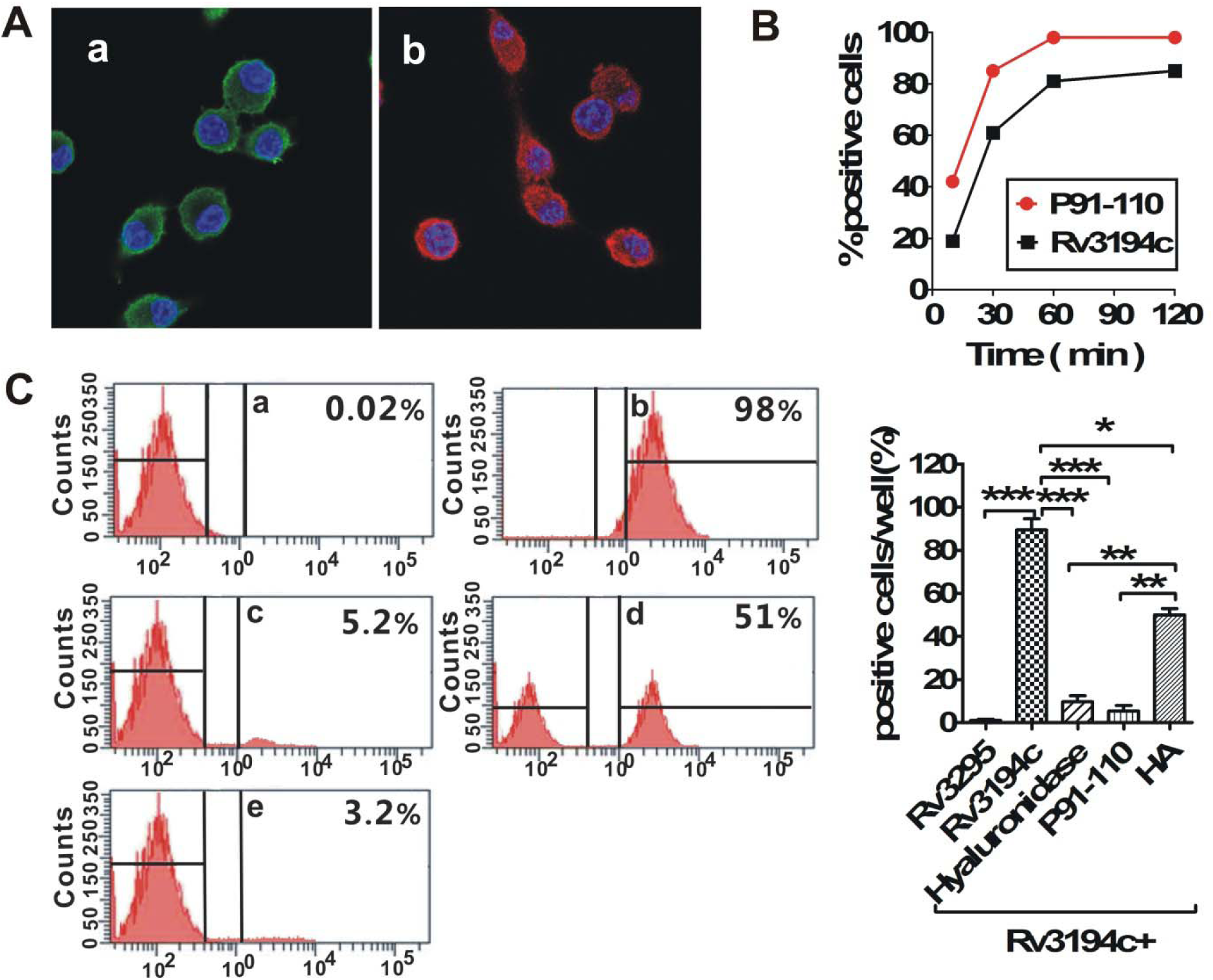
Rv3194c binds to A549 human lung epithelial cells. **A**, The binding of Rv3194c (**a**) and Cy3-P91-110 (**b**) to A549 cells was visualized by confocal microscope. Magnification fold (600×). **B**, Binding kinetics of Cy3-P91-110 and Rv3194c to the A549 cells. Cy3-P91-110-bound cells and Rv3194c-bound cells were identified using FACScan at each time point (30, 60, 90, 120 min). **C**, The percentage of Rv3194c-bound A549 cells were identified by FACScan. Untreated A549 cells bound to Rv3295 as negative control (**a**) and to Rv3194c (**b**). Rv3194c was added to the culture of cells pretreated with hyaluronidase (**c**), HA (**d)** and P91-110 (**e**).

### Rv3194c participates in the attachment of recombinant M. smegmatis to the A549 cells

BLAST analysis revealed that the nucleotide sequence of *Rv3194c* is highly conserved in pathogenic mycobacteria. To clarify the involvement of Rv3194c in the attachment and invasion by the mycobacteria, A549 cells were infected with recombinant *M. smegmatis* (Rv3194c/MS and pMV261/MS) *in vitro*. A549 cells attached by Rv3194c/MS were visualized by confocal laser microscopy, indicating that Rv3194c participates in binding of Rv3194c/MS to A549 cells (**Fig. 5A**). CFU-based analyses showed that bacterial counts of Rv3194c/MS in untreated group significant higher than that in HA or P91-110 pretreated group (**Fig. 5B**). Bacterial counts of Rv3194c/MS in P91-110 pretreated group was significantly lower than that of HA added group (**Fig. 5B**). These results demonstrated that Rv3194c participates in attachment of Rv3194c/MS to A549 cells, and P91-110 treatment of A549 cells almost inhibited Rv3194c/MS-A549 cells interaction and addition of exogenous HA reduced the binding of Rv3194c/MS to A549 cells (**Fig. 5**). We next examined where Rv3194c in Rv3194c/MS was localized using western blotting analyses. The results demonstrated that the Rv3194c expressed in cell lysate, cell wall, cytoplasm and cell membrane (**Fig. 5C**). Rv3194c was detected in cell culture filtrates as well, indicating that the Rv3194c is a secreted protein (**Fig. 5C**).These data demonstrated that the Rv3194c is required when mycobacteria adheres its host cells.

**FIG 5.**
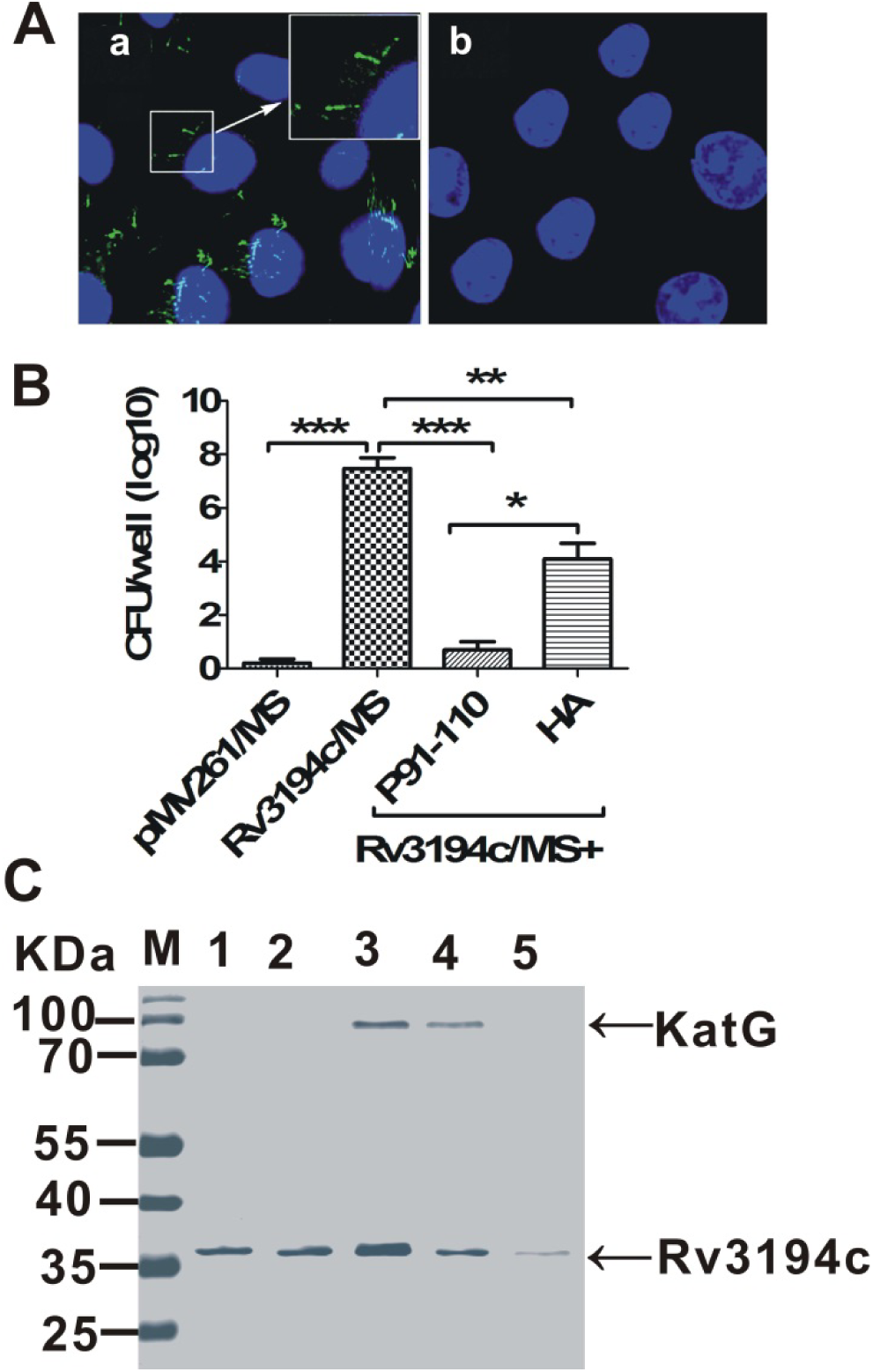
Rv3194c is involved in the recombinant *M. smegmatis* process of attachment and binding to the A549 cells. **A**, The attachment of the Rv3194c/MS (**a**) and pMV261/MS (**b**) to the A549 cells was visualized by confocal microscope. pMV261/MS served as a negative control. **B**: The binding was measured using bacterial count. Untreated A549 cells were infected with pMV261/MS as negative control and with Rv3194c/MS. Rv3194c/MS was added to the culture of cells pretreated with P91-110 and HA. **C**, Subcellular localization of Rv3194c in Rv3194c/MS was detected by western blotting. Lanes: 1, cell wall; 2, cell membrane; 3, whole-cell lysates; 4, cytoplasm; 5, cell culture filtrates. KatG protein serves as a cytoplasm marker.

### Persistence in the lung of the recombinant M. smegmatis expressed Rv3194c

The epithelial cells are target cells for adherence and invasion by mycobacteria (28). Therefore, we expanded our *in vitro* experiments to examine animals *in vivo* to delineate the roles that the Rv3194c play in the mycobacterial/host interaction. To monitor the progression of disease in living animals using bioluminescence imaging, we constructed a reporter strain of *M. smegmatis* including Rv3194c+FFlucRT/MS and FFlucRT/MS (**Fig. 6A**). There was no significant difference among the four groups 1 day after infection (**Fig. 6B**). The bacterial loads were notably higher after infection with Rv3194c+FFlucRT/MS at 4, 7 and 14 days compared to other three groups (**Fig. 6B**). Bacterial loads in the lung tissues of FFlucRT/MS group were completely cleared 14 days after infection (**Fig. 6B**). Furthermore, bacterial loads in the lung tissues of the Rv3194c+FFlucRT/MS plus of P91-110 group were significantly lower than that in Rv3194c+FFlucRT/MS plus of HA group at 7 days and almost cleared at 14 days (**Fig. 6B**). Images of the infected mice were captured at multiple time points after the administration of intraperitoneal luciferin. Imaging mice that had been inoculated with *M. smegmatis* were used to estimate the bioluminescence background level (**Fig. 6C**). The live mice were subjected to *in vivo* imaging, and bioluminescence levels were no significant difference among the four groups 1 day after infection (**Fig. 6C**). Bioluminescence levels in the lung tissues of Rv3194c+FFlucRT/MS group were significantly higher than that in other three groups at 4, 7 and 14 days (**Fig. 6C**). Bioluminescence levels in the lung tissues of Rv3194c+FFlucRT/MS plus of P91-110 group was significantly lower than that in added HA group at 14 days and was close to background level (**Fig. 6C**). Using bioluminescence imaging and bacterial counts confirmed that Rv3194c enhanced the persistence of *M. smegmatis*, and P91-110 more efficiently prevents mice to be infected with Rv3194c+FFlucRT/MS (**Figs. 6B, C**). As expected, there was a strong correlation between the bioluminescence in the thorax of the live mice and CFU in the lungs. These findings demonstrated that Rv3194c serves as a novel adhesin, and P91-110 efficiently suppress the growth of mycobacteria *in vivo* and may be a candidate drug against tuberculosis.

**FIG 6.**
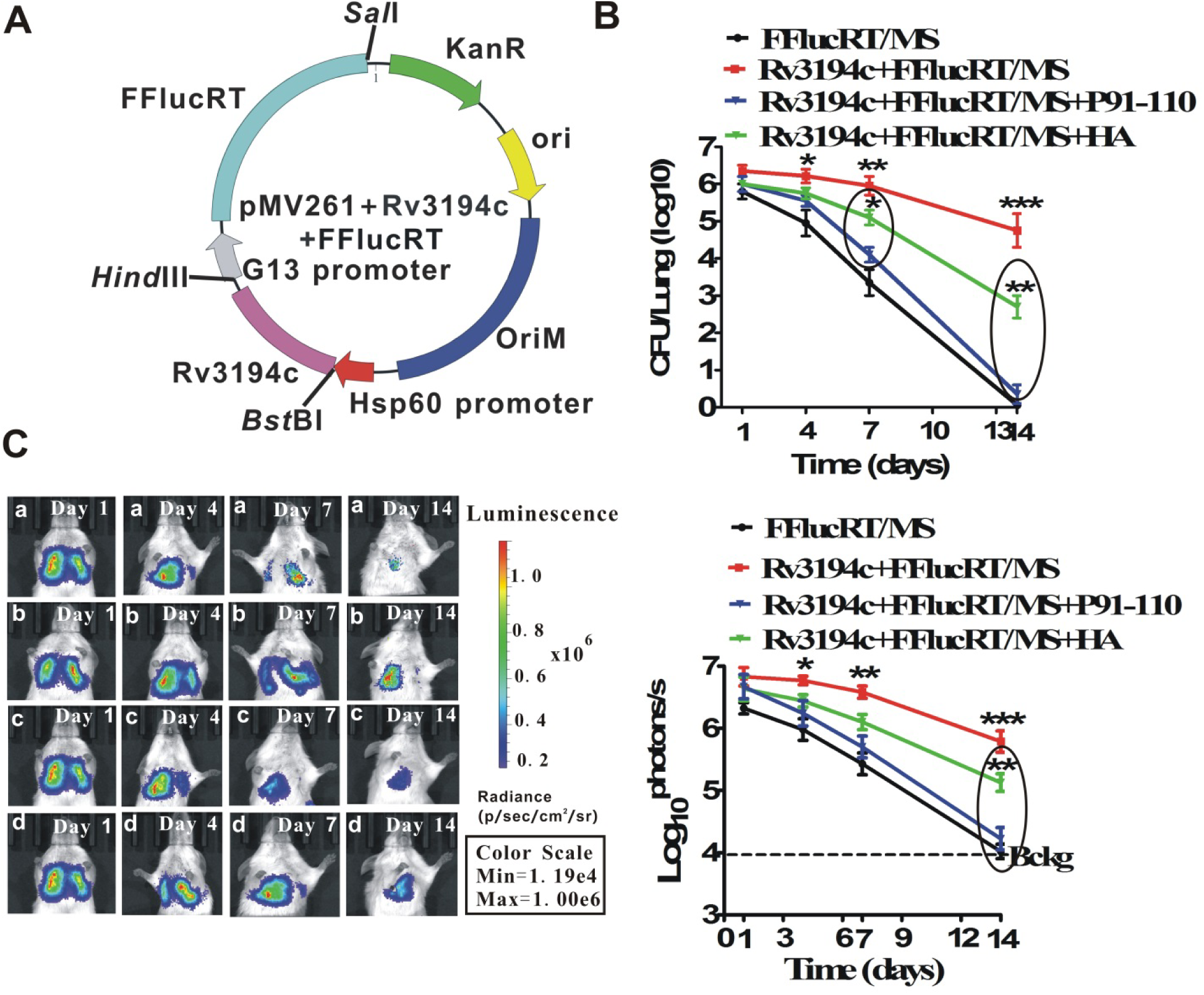
Inhibition of the infection of *in vivo* recombinant *M. smegmatis*. **A**, Construction of the FFlucRT reporter plasmids and strains. **A**, *Rv3194c* and *FFlucRT* containing the G13 promoter were respectively cloned into the *E. coli-mycobacterium* shuttle vector pMV261 at the *Bst*B I-*Hind III* and the *Hind III*-*Sal* I sites. **B and C**, Bacterial loads (**B**) and bioluminescence **(C)** in infected lung tissue from BALB/c mice. The mice was infected with FFlucRT/MS as a negative control (**a**), with Rv3194c+FFlucRT/MS (**b**), with Rv3194c+FFlucRT/MS plus of P91-110 (**c**) and with Rv3194c+FFlucRT/MS plus of HA (**d**) by nasal administration. Bckg, background luminescence (1.0×10^4^ photons/s).

## Discussion

Persistent use of antibiotics will result in death of all susceptible bacteria and generate resistant bacteria through spontaneous mutation in a population (29). Therefore, only those with the mutation can propagate, resulting in the quick spread of resistance in a population. Anti-adhesion therapy represents a potentially promising avenue for the treatment and prevention of tuberculosis in a post-antibiotic era (30). In anti-adhesive therapy, sensitive bacteria are still viable, where resistance to antibiotic therapy has been observed to occur at a much slower rate (31). An anti-adhesion peptide is an efficient strategy to prevent this association, causing the pathogen to be removed by the host, thus preventing disease or treat bacterial infection (32). A surface protein streptococcal antigen (SA) I/II participates in colonization of *Streptococcus mutans* to tooth through binding to salivary receptors adsorbed on the hydroxyapatite matrix of the tooth surface (33). For this area, peptide has been designed, in response to which the inhibition of the connection declines by 65-85% (33). Fuzeon, peptide-based HIV fusion inhibitor, blocks the binding and fusion of viral particles to host cells (34). Peptide-based inhibitors, adhesion analogs as anti-adhesion agents, may be widely successful for anti-adherence therapeutics when applied to specific pathogens. Our works provide evidence that serine protease (Rv3194c) is a multifunctional protein, besides its traditional enzymatic role (26), has evolved to promote the adherence of the bacterium to host cells by its ability to bind HA. The Rv3194c protein belongs to the S16 serine protease family. Analysis of sequence has shown that the B(X_7_)B motif (25) was found in Rv3194c protein, which only exists in pathogenic mycobacteria including *M. tuberculosis, M. africanum* and *M. bovis*. Therefore, the protein is hypothesized to be associated with mycobacterial adherence. To verify this hypothesis, we performed the following studies. Our focus was on the physiological role of the extracellular Rv3194c. In our research, we explored the interaction of Rv3194c with HA using ELISA and surface plasmon resonance (*SPR*) (**Figs. 2A, B**). Utilizing synthetic peptides, we next defined the peptide corresponding to an amino acid sequence of Rv3194c at the 91-110 position (P91-110) is responsible for the specific HA-binding site of Rv3194c (**Fig. 2C**). The higher affinity for binding of Rv3194c or P91-110 to HA was determined using *SPR* analysis (**Fig. 3**). The capability of interaction of Rv3194c with A549 cells was abolished after addition of hyaluronidase or P91-110 to culture of cells, indicating that the surface HA of A549 cells, as the primary receptor, was recognized by P91-110 (**Fig. 4**). Experimental infection *in vitro*, treatment with P91-110 significantly inhibited the mycobacterial binding to the A549 cells compared to that of HA (**Fig. 5B**). Rv3194c in Rv3194c/MS appears on cell wall, cytoplasm, cell membrane and cell filtrates, which further confirms that it is truly an extracellular protein (**Fig. 5C**). This result provides confirmation that Rv3194c was transported out of cytoplasm through cell wall, localized on the outer surface of the mycobacterial cell and released to extracellularly. To our knowledge, this is the first study to demonstrate that a mycobacterium utilizes Rv3194c to attach to its host cells. Given our findings from the *in vitro* experiments, we attempted to therapeutically intervene with infection in a murine model. The treatment of such mice with P91-110 produced a striking decrease of *M. smegmatis* expressing Rv3194c in the lung (**Fig. 6B**). These findings back up the *in vitro* experiments that found that Rv3194c participates in the mycobacterial/lung epithelial cell interaction. To further confirm its role in adhesion, we constructed a reporter strain of *M. smegmatis* which expressed firefly luciferase and Rv3194c to monitor the progression of the disease in the living animals. The bioluminescence imaging in the lung tissue further confirmed that Rv3194c is required for infection, because the persistent ability of recombinant *M. smegmatis* was considerably enhanced compared to that of the parent strain (**Fig. 6C**). In addition, growth and bioluminescence levels of reporter strain that expressed Rv3194c was marked reduced when the mice was treated with exogenous P91-110 (**Figs. 6B, C**). In this study, Rv3194c was identified as a HA-binding adhesion and P91-110 as anti-adhesion agents, adhesion analogs, will be a promising drug against TB.

## Materials and methods

### Cloning, expression and purification

The *Rv3194c* coding sequence without the predicted transmembrane fragment was amplified from *M. tuberculosis H37Rv* genomic DNA using the following primers: the sense primer *Rv3194c*-F: 5’-TAT<u>**CATATG**</u>CCGTTTGTGGCGCTGG-3’, containing an *Nde* I digestible site; the antisense primer *Rv3194c*-R: 5’-GGT<u>**GAATTC**</u>CTAGCAGCTCGGCGTCG-3’, including an *Eco*R I site. PrimeSTAR Max DNA polymerase from TaKaRa Bio (Otsu, Japan) was used to amplify the PCR. The *Rv3194c* amplicon was cloned into a pET28a (+) vector (Novagen, Darmstadt, Germany) that had a 6×His-tag at its N-terminus. The recombinant plasmid pET-28a-*Rv3194c* was then transformed into *E.coli* (BL21) cells. A signal clone was selected and grown in LB media (200 μg/mL ampicillin and 34 μg/mL chloromycetin) at 37 °C overnight. The overnight culture was inoculated into fresh LB medium with kanamycin (50 μg/mL) until their OD_600_ reached 0.6 to 1.0. At that point, they were induced to express the protein by induction with isopropylβ-_D_-1-thiogalactopyranoside (IPTG) at a concentration of 0.5 mM at 23 °C for 12 h. Affinity chromatography using Ni Sepharose 6 Fast Flow (GE Healthcare Life Sciences, Little Chalfont, United Kingdom) was used to purify the protein as previously described (35). The purified protein was centrifuged by a 20 kDa-cut-off membranes made by Millipore (Milford, MA, USA) until the volume was approximately 1 mL. The concentrated protein was then loaded onto superdex75 (10/60) chromatography column (GE Healthcare) pre-equilibrated with 20 mM Tris-HCl buffer at pH 7.4. The desired protein was then eluted by 200 mM NaCl in Tris-HCl buffer at pH 7.4. A flow rate of 1 ml/min was maintained, and 2 ml of each tube was collected. The eluted fractions were collected and analyzed by SDS-PAGE. Finally, Rv3194c was completely identified by mass spectrometry as previously described (36).

### Construction of the FFlucRT reporter strains and polyclonal anti-Rv3194c sera

We used the pMV306DIG13+*FFlucRT* (addgene Plasmid #49998) as a PCR template to amplify the G13 promoter and the *FFlucRT* gene fragment using the following primers: the sense primer: 5’-**<u>AAGCTT</u>**GCAGCCGAACGACCGA-3’ containing a *Hind* III digestible site; the antisense primer: 5’-**<u>GTCGAC</u>**TCTAGATCACAATTTCGA-3’ containing a *Sal* I digestible site. The underlined sections indicate the restriction sites. A *Hind* III-*Sal* I 2.179-kbp fragment containing the G13 promoter sequence and the gene that encodes FFlucRT was cloned into an *E. coli-mycobacterium* shuttle vector pMV261 to produce the plasmid pMV261-*FFlucRT*. To obtain pMV261-*FFlucRT-Rv3194c, Rv3194c* coding sequence including the predicted transmembrane helices was amplified using the following primers: the sense primer *Rv3194c*-F: 5’-**<u>TTCGAA</u>**GTGAATAGGCGGATATTGACC-3’, containing an *Bst*B I digestible site; the antisense primer: *Rv3194c*-R: 5’-**<u>AAGCTT</u>**CTAGCAG CTCGGCGTCG-3’, including a *Hind* III digestible site, and then was cloned into plasmid pMV261-*FFlucRT* by its digestible sites. The rMS (Rv3194c+FFlucRT/MS, FFlucRT/MS, Rv3194c/MS and pMV261/MS) was constructed by the electroporation of *M. smegmatis* with the equivalent vector. The rMS was grown in 7H9 broth at 37 °C until the OD_600_ was 0.8 to 1.0. 1 mg of Rv3194c emulsified in incomplete Freund’s adjuvant was injected multiple times (0, 20 and 40 days) into female rabbits to produce polyclonal anti-Rv3194c sera. Sera were collected as previously described (37). The polyclonal anti-Rv3194c antibody was obtained as previously described (37).

### Preparation of the subcellular fractions of the rMS

To obtain the rMS subcellular fractions, all the subsequent procedures were conducted at 4°C. The rMS cells were resuspended in proteinase inhibitor cocktail, sonicated for 15 min on an ice bath, let to settle for 15 min and again sonicated same as before. The unbroken cells were removed by centrifugation at 3,000 *g* for 5 min. The supernatant and the pellet from supersonic lysate were respectively collected by centrifugation at 27,000 *g* for 1 h. The pellet was resuspended in lysis buffer plus1mM PMSF and centrifuged one time at 27,000 *g* for 1 h. The pellet from this centrifugation was resuspended in 10 mM ammonium bicarbonate. This preparation served as the cell wall fraction. The resultant supernatant (the first and second centrifugations at 27,000 *g*) were pooled and centrifuged at 100,000 *g* for 4 h. After centrifugation, the supernatant and pellet were respectively used as the cytoplasmic and membrane-ribosome fractions. Each fraction was tested with polyclonal antibodies against Rv3194c. The results were detected and recorded using a Li-Cor Odyssey imaging system (Li-Cor Biosciences, USA)

### Detection of the binding of Rv3194c to the HA using enzyme-linked immunosorbent assays

HA, Dextran, Mannose and galactose were respectively immobilized on a 96-well ELISA plate manufactured by Sumitomo (Osaka, Japan) at a concentration of 100 μg/mL in carbonate buffer, pH 7.4, overnight at 4 °C. Following the blockage of the wells with 5% BSA in PBS, 10 μg/mL of Rv3194c in PBS that contained 0.05% Tween 20 (PBS-T) was added to each well and incubated for 1 h at 37 °C. After three washes, anti-Rv3194c antisera diluted 1:1000 in PBS-T was added and incubated for 2 h at 37 °C. After three washes, anti-rabbit IgG antibody conjugated with peroxidase (1:10000) was added and incubated for 1 h at 37 °C. Following three washes, 3’,5,5’-tetramethylbenzidine (TMB), as Chromogenic substrate, was added and incubated for 10 min at 37 °C. A stop solution was utilized to cease the enzymatic chromogenic reaction, and then optical density was read at 630 nm.

### HA-Sepharose Chromatography

100μg of Rv3194c in PBS containing 0.15 M NaCl were applied to HA-Sepharose chromatography (Amersham Biosciences). The unbound proteins were removed by washing the gel with PBS containing 0.15 M NaCl. The bound proteins were eluted by PBS containing 1.5 M NaCl. A flow rate of 1 ml/min was maintained, and 1 ml of each fraction was collected. The fraction was analyzed by Western-blotting.

### Determination of the HA-binding site by ELISA

20-mer of synthetic sequential peptides corresponding to the amino acid sequence of Rv3194c were immobilized on the 96-well ELISA plate (Sumitomo, Osaka, Japan) at a concentration of 10 μg/mL in carbonate buffer, pH 9.6, overnight at 4 °C. After blocking the wells by 5% BSA in PBS, 100 μg/ml of biotinylated HA (Sigma) was added to each well and incubated for 1 h at 37 °C. After washing unbound HA, horseradish peroxidase-conjugated streptavidin was added, and the mixture was incubated for 1 h at 37 °C. After washing free streptavidin, binding was detected by color development with *o*-phenylendiamine dihydrochloride (Wako, Tokyo, Japan), and ELISA units (optical density) were measured at 490 nm.

### Surface Plasmon Resonance Measurements

The interaction of Rv3194c or peptide corresponding to an amino acid sequence of Rv3194c at the 91-110 position (P91-110) with HA was monitored by determining the SPR using a BIAcore 2000 biosensor (Uppsala, Sweden). All the reactions were conducted in binding buffer (10 mM HEPES, 0.005% P20 surfactant, 3 mM EDTA and 150 mM NaCl), at 25 °C. The Rv3194c and P91-110 were immobilized on a BIAcore CM5 sensor chip using an amine coupling kit as described by the manufacturer. The calculation of the association and dissociation rate constants was conducted by nonlinear fitting of the primary sensorgram data using version 3.0 of BIAevaluation software.

### Protein binding assay for the A549 type II human lung epithelial cells

A549 cells were cultured in complete culture media (RPMI 1640 media, 10% fetal bovine serum, 2 mM L-glutamine, 5.5 × 10^−5^ M 2-mercaptoethanol and 25 mM HEPES). 10×10^5^ cells/mL was allotted into individual wells of a 6-well plate (BD Biosciences) and incubated in a humidified atmosphere that contained 5% CO_2_ for 24 h at 37°C. After twice washes with RPMI 1640 media and refilling the volume with 1mL of complete culture media, cells were treated with 100 μg/mL of hyaluronidase, HA or P91-110 for 2 h. The untreated cells and treated cells were respectively incubated with Rv3194c for 1h. After incubation, the wells were washed with RPMI 1640 media three times to remove the free proteins. Then cells were collected by cell scrapers, centrifuged at 600 rpm for 5 min and resuspended in PBS. After washes twice with PBS and centrifugation at 600 rpm for 5 min, cells were fixed by adding 1 ml of 1% paraformaldehyde-PBS. Following washes twice with PBS, the cells were incubated with anti-Rv3194c antibody (1:100) for 2 h. After incubation with primary antibody, cells were washed with PBS for three times and incubated with goat anti-rabbit IgG conjugated with Alexa Fluor 488 (1:10000) for 1h. The cells were analyzed by confocal microscopy and flow cytometry using Cellquest™ software (BD Biosciences, San Jose, CA, USA).

### Experimental Infection in Vitro

Rv3194c/MS was prepared as described above. The Rv3194c/MS were harvested in 7H9 broth at 37 °C until the cells reached an OD_650_ of 0.6. The bacterial suspension was prepared by the conventional method (38). A549 cells were treated with 100 μg/mL of HA or P91-110 dissolved in binding medium (138 mM NaCl, 8.1 mM Na_2_HPO_4_, 1.5 mM KH_2_PO_4_, 2.7 mM KCl, 0.6 mM CaCl_2_, 1 mM MgCl_2_, and 5.5 mM D-glucose) for 2 h. Rv3194c/MS at an MOI of 10:1 was subsequently added to these treated cells or untreated cells. After infection with Rv3194c/MS, the cells were washed three times with RPMI 1640 media, scraped with a rubber policeman and centrifuged. The pellet was then resuspended in 7H9 broth, and the cell suspension were grown on 7H10 agar plates. The colonies were incubated at 37º C for three days and then counted. Confocal microscopy and flow cytometry was used to assess the association of Rv3194c/MS with A549 cells as described in the methods for the protein binding assay.

### Experimental Infection in Vivo

Barrier-bred female 6-12 week BALB/c mice used for the lung infection assays were obtained from the Shanghai Laboratory Animal Center, Chinese Academy of Science (SLACCAS), maintained in specific pathogen-free (SPF) conditions. After the mice were randomly divided into four groups, they were anaesthetized with dry ice before the bacteria were infected. The10^7^ CFUs of rMS (Rv3194c+FFlucRT/MS, FFlucRT/MS) were inoculated the mice by nasal administration. In addition, the 10^7^ CFUs of Rv3194c+FFlucRT/MS were mixed with100 μg of HA or P91-110, and the mixture was also inoculated other mice by nasal administration. At 1, 4, 7 and 14 days post infection, bioluminescence imaging and bacterial counts in the lung tissue were performed to track the disease progression. The bioluminescence [photons/s/cm^2^/steradian (sr)] was assessed from the living animals using an IVISw Spectrum system. The mice were intraperitoneally injected with 500 mg/kg body weight of D-luciferin dissolved in sterile D-PBS containing ketamine before the bioluminescent imaging. The mice were confined to a large airtight box to ensure safe conditions and placed into the imaging chamber of the IVISw Spectrum imaging system with the stage that had been heated to 37 °C. The bioluminescence within specific regions of the individual mice was also quantified by the ROI tool in the Living Image software program (set as photons/s). The *in vivo* imaging was immediately performed at specific time points. The animals were culled while still under anesthesia by cervical dislocation, and the lungs were aseptically removed. The entire left lobe of the lung was homogenized with a set of motor and pestle, and then was plated on 7H10 agar plates with 50 μg/mL kanamycin. After 3 days of incubation at 37 °C, the CFU of the rMs was calculated.

### Statistical analysis

Each experiment was repeated in triplicate. A two-tailed Student’s t test was used to determine the statistical significance, and Prism software (version 5.0; GraphPad, San Diego, CA, USA) was used for these analyses. Data that appeared to be statistically significant were analyzed using an analysis of variance (ANOVA) to compare the means of multiple groups. The means were considered to be significant if the *P* values were less than 0.05 (0.01< \**P* < 0.05, \*\**P* < 0.01 and \*\*\**P* < 0.001).

## Acknowledgments

This study was supported by the National Natural Science Foundation of China (No. 31901930), Educational Department Project (Category A) of Fujian Province (No. JT180080) and the scientific research innovation program “Xiyuanjiang River Scholarship” of College of Life Sciences of Fujian Normal University.

## Author contributions

D.Z., C.X. and D.L. conceived and designed the experiments, analyzed the data. D.Z., C.X. and D.L. performed the experiments. D.Z. wrote the manuscript.

## Ethical approval

The animal use protocol listed has been reviewed and approved by the Animal Ethical and Welfare Committee (AEWC) of Fujian Normal University (FJNU) (No. of Animal use permit: SYXK: 2015-0004, Approval No. IACUC-20190033).

## Data availability statement

All data generated or analyzed during this study are included in this article.

## References

1. Bloom BR, Salomon JA. 2005. Enlightened self-interest and the control of tuberculosis. N Engl J Med 353:1057–1059.

2. Ribet D, Cossart P. 2015. How bacterial pathogens colonize their hosts and invade deeper tissues. Microbes Infect 17:173–183.

3. Mehta PK, Karls RK, White EH, Ades EW, Quinn FD. 2006. Entry and intracellular replication of *Mycobacterium tuberculosis* in cultured human microvascular endothelial cells. Microb Pathog 41:119–124.

4. Stones DH, Krachler AM. 2015. Fatal attraction: how bacterial adhesins affect host signaling and what we can learn from them. Int J Mol Sci 16:2626–2640.

5. Boland T, Latour RA, Stutzenberger FJ. Molecular basis of bacterial adhesion. In: An YH, Friedman RJ editors. Handbook Handbook of bacterial adhesion: principles, methods, and applications. Totowa: Humana Press Inc; 2000. p. 29–41.

6. Asadi A, Razavi S, Talebi M, Gholami M. 2019. A review on anti-adhesion therapies of bacterial diseases. Infection 47:13–23.

7. Ramsugit S, Pillay M. 2014. *Mycobacterium tuberculosis* pili promote adhesion to and invasion of THP-1 macrophages. Jpn J Infect Dis 67:476–478.

8. Ramsugit S, Pillay B, Pillay M. 2016. Evaluation of the role of *Mycobacterium tuberculosis* pili (MTP) as an adhesin, invasin, and cytokine inducer of epithelial cells. Braz J Infect Dis 20:160–165.

9. Brennan MJ, Delogu G, Chen Y, Bardarov S, Kriakov J, Alavi M, Jacobs WR. 2001. Evidence that mycobacterial PE-PGRS proteins are cell surface constituents that influence interactions with other cells. Infect Immun 69:7326–7333.

10. Kinhikar AG, Verma I, Chandra D, Singh KK, Weldingh K, Andersen P. 2000. Potential role for ESAT6 in dissemination of *M. tuberculosis* via human lung epithelial cells. Mol Microbio 75:92–106.

11. Hickey TB, Ziltener HJ, Speert DP, Stokes RW. 2010. *Mycobacterium tuberculosis* employs Cpn60.2 as an adhesin that binds CD43 on the macrophage surface. Cell Microbiol 12:1634–1647.

12. Chitale S, Ehrt S, Kawamura I, Fujimura T, Shimono N, Anand N. 2001. Recombinant *Mycobacterium tuberculosis* protein associated with mammalian cell entry. Cell Microbiol 3:247–254.

13. Kumar S, Puniya BL, Parween S, Nahar P, Ramachandran S. 2013. Identification of novel adhesins of *M. tuberculosis* H37Rv using integrated approach of multiple computational algorithms and experimental analysis. PLoS One 8:e69790.

14. Kinhikar AG, Vargas D, Li H, Mahaffey SB, Hinds L, Belisle JT, Laal S. 2006. Mycobacterium tuberculosis malate synthase is a laminin-binding adhesin. Mol Microbiol 60:999–1013.

15. Espitia C, Laclette JP, Mondragón-Palomino M, Amador A, Campuzano J, Martens A, Singh M, Cicero R, Zhang Y, Moreno C. 1999. The PE-PGRS glycine-rich proteins of *Mycobacterium tuberculosis*: a new family of fibronectin-binding proteins? Microbiology 145:3487–3495.

16. Abou-Zeid C, Ratliff TL, Wiker HG, Harboe M, Bennedsen J, Rook GA. 1988. Characterization of fibronectin-binding antigens released by *Mycobacterium tuberculosis* and *Mycobacterium bovis BCG*. Infect Immun; 56:3046–3051.

17. Xolalpa W, Vallecillo AJ, Lara M, Mendoza-Hernandez G, Comini M, Spallek R, Singh M, Espitia C. 2007. Identification of novel bacterial plasminogen-binding proteins in the human pathogen *Mycobacterium tuberculosis*. Proteomics 7:3332–3341.

18. Esparza M, Palomares B, García T, Espinosa P, Zenteno E, Mancilla R. 2015. PstS-1, the 38-kDa *Mycobacterium tuberculosis* glycoprotein, is an adhesin, which binds the macrophage mannose receptor and promotes phagocytosis. Scand J Immunol 81:46–55.

19. Diaz-Silvestre H, Espinosa-Cueto P, Sanchez-Gonzalez A, Pereira-Suarez AL, Bernal-Fernandez G, Espitia C, Mancilla R. 2005. The 19-kDa antigen of *Mycobacterium tuberculosis* is a major adhesin that binds the mannose receptor of THP-1 monocytic cells and promotes phagocytosis of mycobacteria. Microb Pathog 39:97–107.

20. Aoki K, Matsumoto S, Hirayama Y, Wada T, Ozeki Y, Niki M, Domenech P, Umemori K, Yamamoto S, Mineda A, Matsumoto M, Kobayashi K. 2004. Extracellular mycobacterial DNA-binding protein 1 participates in mycobacterium-lung epithelial cell interaction through hyaluronic acid. J Biol Chem 279:39798–397806.

21. Pethe K, Aumercier M, Fort E, Gatot C, Locht C, Menozzi FD. 2000. Characterization of the heparin-binding site of the mycobacterial heparin-binding hemagglutininadhesin. J Biol Chem 275: 14273–14280.

22. Russell DG. 2011. Mycobacterium tuberculosis and the intimate discourse of a chronic infection. Immunol Rev 240:252–268.

23. Krachler AM, Orth K. 2013. Made to stick: anti-adhesion therapy for bacterial infections. Microbe 8:286–290.

24. Smith I. 2003. Mycobacterium tuberculosis pathogenesis and molecular determinants of virulence. Clin Microbiol Rev 16:463–496.

25. Yang B, Yang BL, Savani RC, Turley EA. 1994. Identification of a common hyaluronan binding motif in the hyaluronan binding proteins RHAMM, CD44 and link protein. EMBO J 13:286–296.

26. Li H, Dang G, Liu H, Wang Z, Cui Z, Song N, Chen L, Liu S. 2019. Characterization of a novel *Mycobacterium tuberculosis* serine protease (Rv3194c) activity and pathogenicity. Tuberculosis 119:101880.

27. Forteza R, Lieb T, Aoki T, Savani RC, Conner GE, Salathe M. 2001. Hyaluronan serves a novel role in airway mucosal host defense. FASEB J 15:2179e86.

28. Ramsugit S, Pillay M. 2016. Identification of *Mycobacterium tuberculosis* adherence-mediating components: a review of key methods to confirm adhesin function. Iran J Basic Med Sci 19:579–584.

29. Ofek I, Hasty DL, Sharon N. 2003. Anti-adhesion therapy of bacterial diseases: prospects and problems. FEMS Immunol Med Microbiol 38:181–191.

30. Krachler AM, Orth K. 2013. Targeting the bacteria-host interface: strategies in anti-adhesion therapy. Virulence 4:284–94.

31. Sharon N. 2006. Carbohydrates as future anti-adhesion drugs for infectious diseases. Biochim Biophys Acta (BBA) Gen Subj 1760:527–37.

32. Asadi A, Razavi S, Talebi M, Gholami M. 2019. A review on anti-adhesion therapies of bacterial diseases. Infection 47:13–23.

33. Munro GH, Evans P, Todryk S, Buckett P, Kelly CG, Lehner T. 1993. A protein fragment of streptococcal cell surface antigen I/II which prevents adhesion of *Streptococcus mutans*. Infect Immun 61:4590–4598.

34. Lalezari JP, Henry K, O’hearn M, Montaner JS, Piliero PJ, Trottier B. 2003. Enfuvirtide, an HIV-1 fusion inhibitor, for drug resistant HIV infection in North and South America. N Engl J Med 348:2175–2185.

35. Dang G, Cao J, Cui Y, Song N, Chen L, Pang H, Liu S. 2016. Characterization of Rv0888, a novel extracellular nuclease from *Mycobacterium tuberculosis*. Sci Rep 6:19033.

36. Yuan X, Chen L, Deng X, Cao J, Yu S, Quankai W, Pang H, Liu S. 2013. Characterization of Rv0394c gene encoding hyaluronidase and chondrosulfatase from *Mycobacterium tuberculosis*. Tuberculosis 93:296–300.

37. Vizcaíno C, Patarroyo ME, Patarroyo MA. 2010. Computational prediction and experimental assessment of secreted/surface proteins from *Mycobacterium tuberculosis* H37Rv. PLoS Comput Biol 6:e1000824.

38. Hoal-van Helden EG, Hon D, Lewis LA, Beyers N, van Helden PD. 2001. Mycobacterial growth in human macrophages: variation according to donor, inoculum and bacterial strain. Cell Biol Int 25:71–81.

